# BZR1 promotes pluripotency acquisition and callus development through direct regulation of ARF7 and ARF19

**DOI:** 10.1101/2024.03.22.586258

**Authors:** E Ebstrup, T Ammitsøe, N Blanco-Touriñán, J Hansen, CS Hardtke, E Rodriguez, M Petersen

## Abstract

Plants have the remarkable ability to regenerate whole organisms through formation of pluripotent cell masses from somatic cells. Cellular programs leading to fate change of somatic to pluripotent cells resembles lateral root (LR) formation and both are chiefly regulated by auxin. Brassinosteroid signalling also plays an important role during LR formation but little is known about the direct link between auxin and brassinosteroid components, such as BZR1 and BES1, in relation to pluripotency acquisition. Here we show that gain-of-function mutants *bzr1-D* and *bes1-D* exhibit altered callus formation, yet disruption of these transcription factors does not produce major changes to callus formation or *de novo organogenesis*. Moreover, our data reveals that BZR1 displays enhanced expression in callus tissue and directly binds to the promoters of ARF7 and ARF19, two master pluripotency regulators, leading to their enhanced transcription. Remarkably, we see abrogation of callus formation in *bzr1-D* upon disruption of ARF7 and ARF19, emphasizing that BZR1 callus phenotype is dependent on these two auxin signalling components. In conclusion, we depict a link between ARF7, ARF19 and BZR1 in the promotion of pluripotency acquisition, portraying BZR1 as a major supporting factor in callus formation.

**IMPORTANT:**

- Manuscripts submitted to Review Commons are peer reviewed in a journal-agnostic way.
- Upon transfer of the peer reviewed preprint to a journal, the referee reports will be available in full to the handling editor.
- The identity of the referees will NOT be communicated to the authors unless the reviewers choose to sign their report.
- The identity of the referee will be confidentially disclosed to any affiliate journals to which the manuscript is transferred.

**GUIDELINES:**

- For reviewers: https://www.reviewcommons.org/reviewers
- For authors: https://www.reviewcommons.org/authors

**CONTACT:** The Review Commons office can be contacted directly at

## Introduction

Plants possess extraordinary plasticity, being able to naturally dedifferentiate somatic cells back into pluripotency allowing control of differentiation and self-renewal through transcriptional, post-transcriptional, and epigenetic modifications (Li and Belmonte, 2017). The steps necessary for somatic cells to achieve pluripotency and redifferentiation for *de novo* organogenesis can be studied *in vitro*, through a process referred to as callus formation (Bustillo-Avendaño et al., 2018; Ikeuchi et al., 2019, 2017, 2016). Callus formation can be induced through tissue culture in callus-inducing media (CIM; high auxin – low cytokinin ratio) (Valvekens et al., 1988), which promotes an overdrive of hormone developmental programs to unlock pluripotency (Gordon et al., 2007a; Sang et al., 2018). In particular, under CIM treatment, Xylem Pole Pericycle (XPP) cells undergo cell fate change, re-enter the cell cycle and display increased gene expression of quiescent center (QC) markers such as *WUSCHEL-RELATED-HOMEOBOX5* (*WOX5*), *SHORTROOT* (*SHR*), and *SCARECROW* (*SCR*) (Ikeuchi et al., 2019; Sugimoto et al., 2010) – a process identical to lateral root (LR) initiation (Che et al., 2007; Iwase et al., 2017; Sugimoto et al., 2010). Both LR and callus formation are regulated by an auxin responsive signaling node composed by the repressor INDOLE-ACETIC-ACID14 (IAA14)/SOLITARY-ROOT (SLR), transcriptional activators; AUXIN RESPONSIVE FACTOR7 (ARF7) and AUXIN RESPONSIVE FACTOR19 (ARF19), and their direct downstream targets LATERAL ORGAN BOUNDARIES DOMAINS (LBDs) in such a way that auxin induces the degradation of IAA14 to unlock ARF7 and ARF19 transcriptional output (Fukaki et al., 2002; Goh et al., 2012; Okushima, et al., 2007; Okushima et al., 2005). The relevance of this signaling node to both callus and LR formation is patented by observations that the double knockout (KO) *arf7-1 arf19-2* or the gain of function mutant *slr* do not make callus or LRs, and overexpressed *LBDs* form ectopic callus (Fan et al., 2012; Iwase et al., 2011). After callus formation, *in vitro* shoot regeneration can be achieved by transferring callus masses to shoot-inducing media (SIM; high cytokinin – low auxin); upon culturing callus tissue in SIM, cells undergo a novel cell fate change, from a root meristem-like structure to forming shoot apical meristems (SAM), with expression of SAM markers like *WUSCHEL* (*WUS*), *CLAVATA3* (*CLV3*) etc., which enables the formation of shoots (Gordon et al., 2007b; Su et al., 2009).

While the role of auxin and cytokinin signalling during callus formation and *de novo* organongenesis has been well established, how other hormone signalling participate in this process is comparably less understood. For instance, the steroid hormone brassinosteroids (BR) are known to be important for the regulation of a range of developmental aspects both above and below ground (e.g. (Bajguz et al., 2020; Hacham et al., 2011; Lee et al., 2019; Oh et al., 2014; Planas-Riverola et al., 2019; Saito et al., 2018)), among them LR formation (Bao et al., 2004; Rovere et al., 2022). BR signaling in the root is mainly required for appropriate cell elongation and cellular anisotropy, and for restricting the extent of formative divisions( Fridman et al., 2021; Graeff et al., 2020; Kang et al., 2017) and has been suggested to contribute to properly intergrated sieve element differentiation (Graeff et al., 2020). BR perception by the leucine-rich-repeat receptor kinase, BRASSINOSTEROID INSENSITIVE 1 (BRI1) leads to the inactivation of the shaggy-like protein kinase BRASSINOSTEROID INSENSITIVE 2 (BIN2) and the dephosphorylation of the BES1/BZR1 BR transcription factor (TF) family, consisting of BRASSINAZOLE RESISTANT 1 (BZR1) and BRI1-EMS-SUPPRESSOR 1 (BES1) and the four BES1/BZR1 Homologs (BEH1-4), which unlock BR responsive genes in the nucleus (Kim et al., 2023, 2009; Manghwar et al., 2022; Ryu et al., 2007; Wang et al., 2002; Yin et al., 2002). The importance of the phosphorylation status to BZR1 and BES1 function have been established by isolation of dominant mutant alleles of both TFs with a point mutation, preventing BIN2 mediated phosphorylation and consequent turnover of BZR1 and BES1 (Wang et al., 2002; Yin et al., 2002). Interestingly, while BZR1 and BES1 share 88% homology in sequence, contain similar domains, (He et al., 2005; Nolan et al., 2020; Sun et al., 2011; Yu et al., 2011) and bind to E-box (CANNTG) sites in the promoter of target genes, the dominant mutants *bzr1-D* and *bes1-D* can exhibit different phenotypes– suggesting two different biological functions concerning plant development (Wang et al., 2002; Yin et al., 2002). Further research proposed that BES1 activates BR biosynthesis and signaling involved in plant growth (Yu et al., 2011), whereas BZR1 could function as a transcriptional repressor, binding directly to promoters of BR components to reduce BR levels (He et al., 2005). In addition to having a role as a transcriptional repressor, BZR1 has also been linked to the promotion of growth, downstream of BR components (He et al., 2005). Contrarily, BZR1 has been suggested to inhibit organ boundary formation, an important process in plant organogenesis, through the repression of *CUP-SHAPED COTYLEDON* (*CUC*) genes in the primordium cells located in the shoot apical meristem (Gendron et al., 2012). Plants exhibiting enhanced BR responses, such as *bes1-D,* display a decrease in procambial cell layers within the hypocotyl, indicating BES1-dependent promotion of xylem differentiation; notably this was not seen in *bzr1-D* (Kondo et al., 2014). The *bes1-D* mutant has also been shown to negatively affect meristem size and growth due to a premature exit of the cell cycle in the early differentiation of columella stem cells (CSC) (González-García et al., 2011). Additionally, BES1 has been shown to control key processes of cell division and cell elongation, for instance, promoting differentiation through *WOX5* expression in QC cells and CSCs (González et al., 2011; Lee et al., 2015) and through repressing of BRAVO activity in the QC zone in the root (Lozano-elena et al., 2018; Vilarrasa-Blasi et al., 2014a).

The fact that auxin typically serves as the primary driving force in root development highlights an interconnected link between BR and auxin. These two hormones are known to share several target genes and have overlapping activities notably in the promotion of LR formation (Bao et al., 2004; Chaiwanon and Wang, 2015; Kim et al., 2007; Li et al., 2005; Mazzoni-putman et al., 2021). Auxin has been shown to promote BZR1’s nuclear accumulation and direct binding and transcriptional control of ARF7 during hypocotyl elongation (Yu et al., 2023), thus establishing a direct molecular link between BZR1 and auxin signaling pathway (Zhou et al., 2013). Similarly, BIN2 has been shown to be a positive regulator of LR organogenesis (Cho et al., 2014) and callus formation (Lee and Seo, 2017) through direct regulation of ARF7 and ARF19. Moreover, a recent study has shown BR to both promote auxin levels and repress the transcriptional output of auxin signaling in established meristems as well as *de novo*-formed meristems (Ackerman-Lavert et al., 2021). Although the interplay between auxin and BR in root development has been linked, the interchange and underlying mechanisms are still not fully understood.

In this study, we investigate the involvement of BZR1 and BES1 transcription factors in the processes of callus formation and *de novo* organogenesis. Specifically, we discovered that BZR1 expression is enhanced in root explants cultured in CIM. Consistent with this, plants expressing the dominant form of BZR1, *bzr1-D,* exhibit amplified callus masses and an increase in the abundance of *ARF7* and *ARF19* transcripts. Furthermore, we identified that BZR1 directly targets the promoters of *ARF7* and *ARF19*, which are crucial for the dedifferentiation of somatic cells into pluripotent cells in *bzr1-D*, revealing an important contribution of brassinosteroid signaling components to callus formation.

## Results and discussion

### Stabilization of BZR1 and BES1 have different outcomes during pluripotency acquisition and de novo shoot formation

Given the interesting proposition of a direct link between auxin and different BR signaling components like BZR1 during developmental changes (e.g. hypocotyl elongation), we wondered if this could also be observed in *bzr1-D* and *bes1-D* during pluripotency acquisition and callus formation. To test this, we cultivated root explants of *bzr1-D* and *bes1-D* on CIM media for 21 days and evaluated their performance during callus formation (Figure 1A). Surprisingly, *bzr1-D* exhibited a significant increase in callus mass compared to the wild type (WT), while *bes1-D* produced slightly lower callus mass compared to WT (Figure 1A-B). The phenotype observed for *bzr1-D* after 21-day CIM, coincides with previous studies showing stimulation of LR formation upon the addition of low levels of exogenous BR (Rovere et al., 2022) and an increase in LR formation by overexpression of *BZR1* in rice (Hou et al., 2022). The decrease in callus mass seen in *bes1-D* could be explained by issues during the cell cycle -and premature differentiation of stem cells, which have been previously reported for CSCs (Vilarrasa-Blasi et al., 2014b).

**Figure 1.**
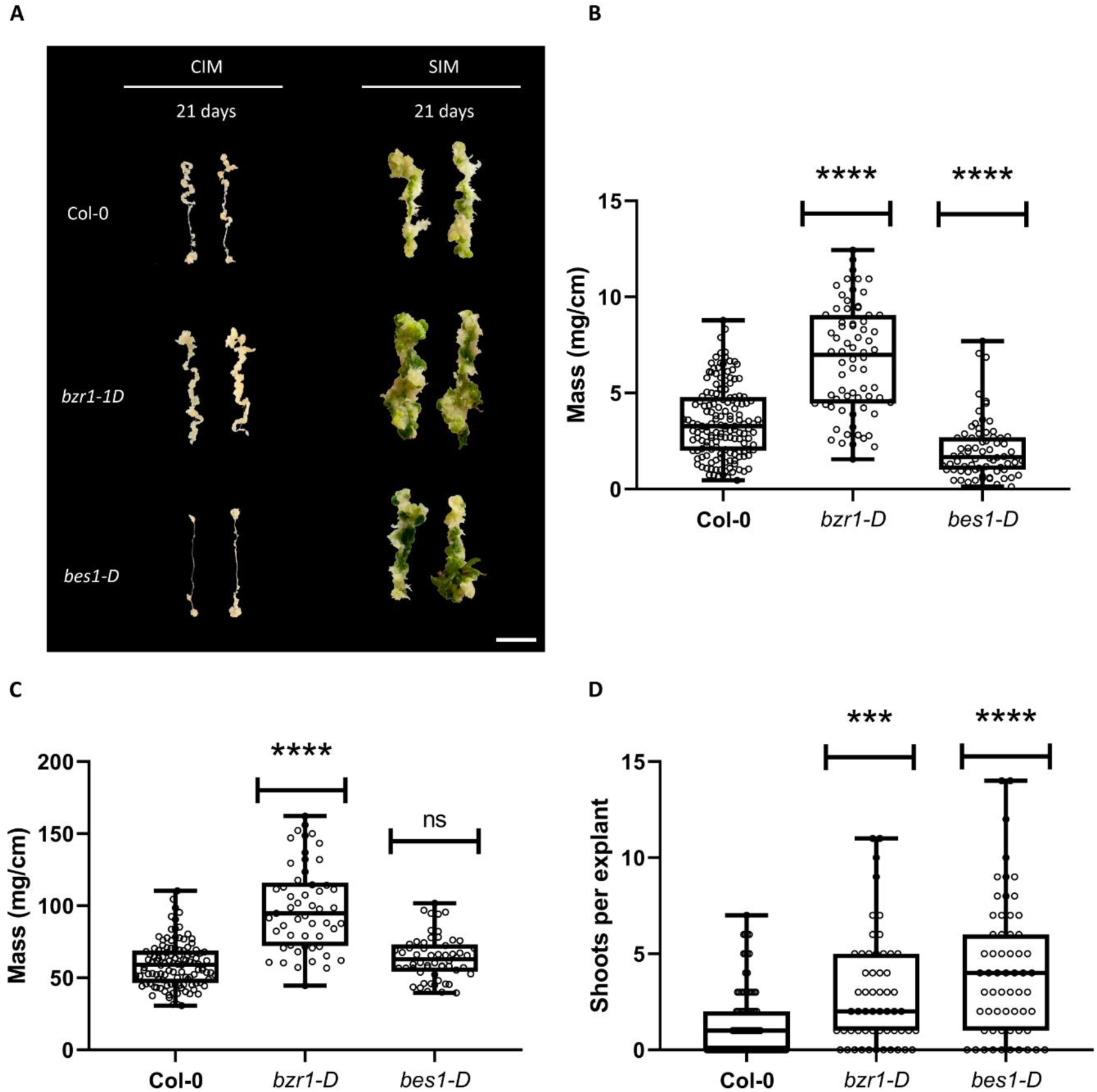
Stabilization of BES1 and BZR1 promote different outcomes during pluripotency acquisition and shot formation. A) Representative images of callus forming root explants WT, bzr1-D, and bes1-D from 21-day CIM or 21-day SIM (21- day CIM + 21-day SIM); scale bar: 1 cm. B) Fresh weight of 21-day root explants on CIM. C) Fresh weight of 21-day-explants on SIM. D) Shoots counted per callus explant after 21 days on SIM. Boxplots: centerlines show the median; the box limits the 25^th^ and 75^th^ percentiles; whiskers extend to the minimum and maximum. Significance was calculated viaKruskal-Wallis non-parametric test (P<0,05) . Asterisks represent the significant difference from WT, (***P<0.001, ****P<0.0001). Experiments were performed in three biological replicates with total n=70-170 (B), n=53-113 (C,D)

Next, we wanted to explore if the differences in callus output between *bes1-D* and *bzr1-D* were also observable during *de novo* shoot formation after CIM treatment. To examine this, we transferred 21-day-old CIM-grown explants to SIM and followed them for another 21 days (Figure 1A). Interestingly culturing *bes1-D* on SIM leads to improvement in callus formation, being able to catch up to WT levels (Figure 1C). We also noticed the shoot formation was significantly higher in *bes1-D* compared to WT (Figure 1D). As for *bzr1-D*, explants from this genotype still displayed a significantly higher amount of callus mass than the WT after SIM incubation. Similar to *bes1-D*, *bzr1-D* also displayed a higher number of shoots compared to WT (Figure 1D). BR has previously been associated with the repression of organ boundary formation important for the shoot apical meristem (Gendron et al., 2012), thus in combination, these results suggest that stabilization and expansion of BES1/BZR1 facilitates and enlarges shoot meristematic area.

### BZR1 and BES1 are not crucial for callus and shoot growth

Intrigued by our results regarding the dominant versions of BZR1 and BES1, we next wanted to examine the callus-related phenotypes of the loss-of-function mutants of these genes. A *BES1* T-DNA line was available from Arabidopsis seed stock center (Alonso et al., 2003) but none was available for *BZR1* and we thus generated them by Crispr-CAS9 gene editing. We isolated two knockout lines of *BZR1*, *bzr1-c1* and *bzr1-c2*, one with an 1bp insertion and the other with a deletion of a single nucleotide (at position 144 and 143 of the *BZR1 cds)* respectively, both resulting in a frameshift mutation disrupting the gene (Figure S1). The three mutants were cultivated on CIM for a period of 21 days as illustrated in Figure 2A. Interestingly, *bes1-ko* and *bzr1-c1* showed no significant difference, whereas *bzr1-c2* displayed increased mass, compared to the WT, however no statistical difference was found between the lines (Figure 2B). This indicates that the absence of these two transcription factors does not negatively impact callus formation. As in Figure 1, we examined the effect on the mutants when transitioning from CIM to SIM. For *bes1-ko, bzr1-c1* and *bzr1-c2* no significant difference was found between the lines or compared to WT in terms of *de novo organogenesis* In line with the results depicted in Figure 2B, no significant differences were found between the mutants and WT in terms of *de novo* organogenesis (Figure 2C-D).

**Figure 2.**
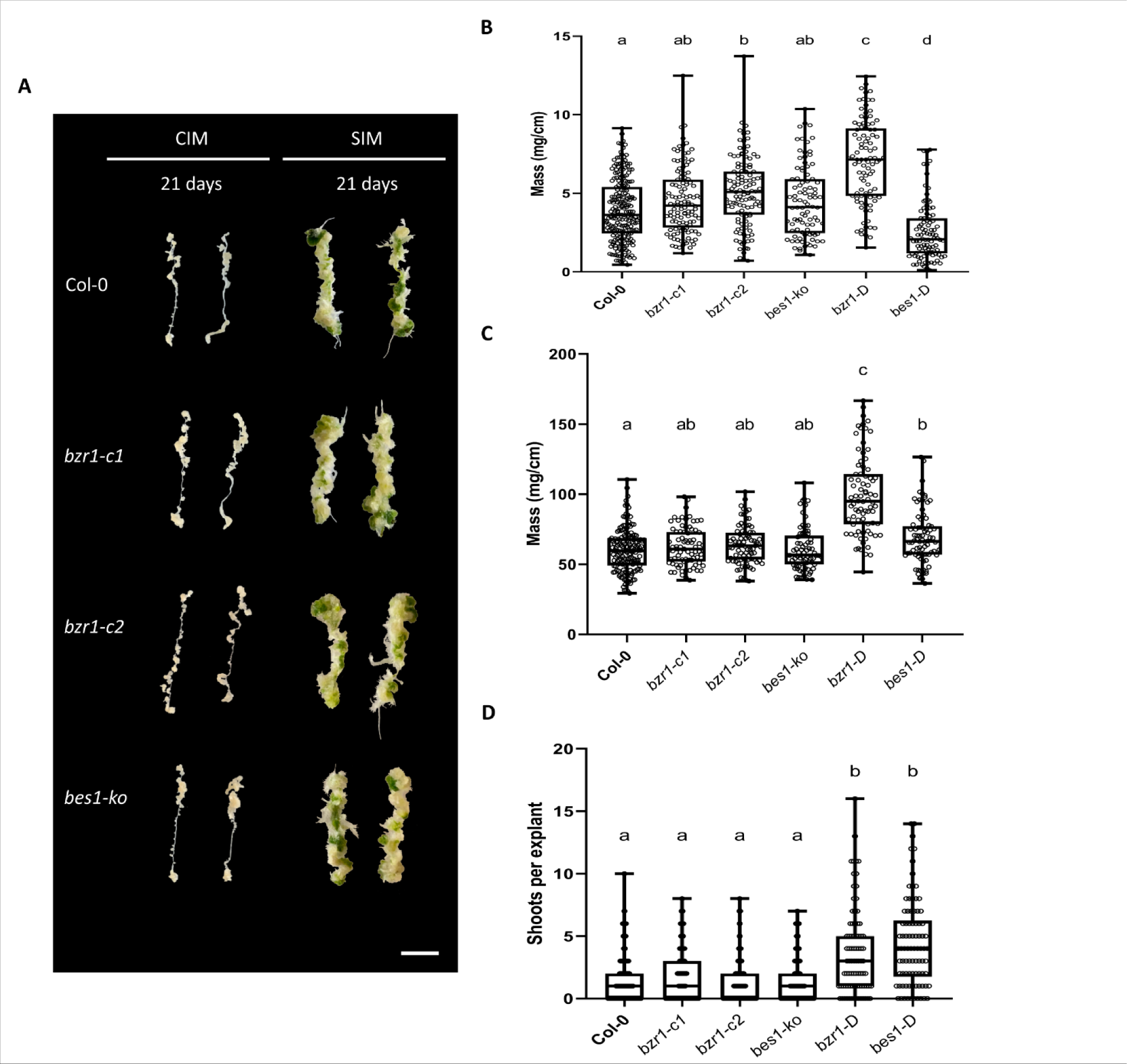
BZR1 and BES1 are dispensable for callus formation. A) Representative images of callus forming root explants WT, bzr1-c1, bzr1-c2 and bes1-KO from 21-day CIM or 21-day SIM (21-day CIM + 21-day SIM); scale bar: 1 cm. B) Fresh weight of 21-day root explants on CIM. C) Fresh weight of 21-day explants on SIM. D) Shoots counted per callus explant after 21-days on SIM. Boxplots: centerlines show the median; the box limits the 25^th^ and 75^th^ percentiles; whiskers extend to the minimum and maximum. Significance was calculated via Kruskal-Wallis non-parametric test with multiple comparisons calculated via Dunn’s test post-hoc (P<0,05). Lower case letters indicates significant difference between samples.. Experiments were performed in three biological replicates with total n=93-204 (B), n=75-159 (C,D).

Our findings indicate that the absence of BZR1 and BES1 do not produce a significant effect on callus or *de novo* shoot formation. However, it has been recently shown that that *BES1* RNAi explants have increased levels of shoots and suggested both BES1 and BZR1 to be negative regulators of shoot formation (Wu et al., 2022). These differences could be explained by the possibility that the RNAi construct used by Wu and co-workers may to some degree also target *BZR1* and even potentially their 4 other homologues (*BEH1-4*), (Yin et al., 2005) which would suggest redundancy in the BES/BZR TF family. Precisely, this redundancy has been observed during vegetal growth where only *BES/BZR* sextuplet mutant displays major developmental defects, resembling the *bri-t* BR insensitive mutant (L.-G. Chen et al., 2019). Likewise, overexpression of each member of the *BZR/BES* family could rescue the *bri1-5* mutant phenotype which also supports the notion of a high degree of redundancy among this family of TF (Kim et al., 2023). It should be noted that, despite the obvious redundancy among the BES/BZR1 family, each member of the family might still differ in their regulation of BR independent genes. For instance, the introduction of mutations to Proline234, causing the dominant mutation in *bes1-D* and *bzr1-D,* does not yield similar developmental phenotypes across the family, suggesting specific differences in this TF family (Kim et al., 2023). Thus, there could be redundancy in the BES/BZR TF family during callus and shoot formation that masks the resulting phenotype of the *bes-ko* and *bzr1-c1/2* lines.

### CIM and SIM promote different fates for BES1 and BZR1 expression

To investigate the expression patterns of *BZR1* and *BES1* in relation to the observed phenotypes in Figure 1, we utilized transgenic lines expressing the translational reporter *pBES1::BES1-Venus* or *pBZR1::BZR1-Venus* (W. Chen et al., 2019) and examined their fluorescence intensity by confocal microscopy upon culturing in CIM . For this purpose, 7-day-old root explants were treated with either 6-days in liquid CIM or 6-days in liquid ½ MS plus solvents. Under ½ MS; BES1-YFP displayed nuclear localization in the root tip (Figure 3A). However, this signal decreased significantly upon CIM treatment (Figure 3A). In mature tissue, no nuclear signal of BES1 was detected on either MS or CIM (Figure 3A). The expression pattern observed with BZR1-YFP showed nuclear localization at the root tip (Figure 3B). Contrary to BES1-YFP, the signal of BZR1-YFP increased in both the nucleus and the cytoplasm following CIM treatment (Figure 3B). Notably, cells surrounding the xylem poles exhibited a strong nuclear signal of BZR1-YFP (Figure 3B). These results are overall in agreement with the phenotypes observed in Figure 1, *i.e.* BZR1 signal appeared to increase upon CIM treatment, while the BES1 signal is reduced. The lower expression of *BES1* could be explained by the fact that callus induction is not commonly associated with BR synthesis, as BES1 has been shown to positively regulate this process (Yu et al., 2011) while BZR1 acts on more downstream responsive genes (He *et al*., 2005). The increased BZR1 signal could stem from the high levels of auxin in the CIM treatment, as a previous study has shown induced nuclear localization of BZR1 in the hypocotyl, under auxin-induced elongation (Yu et al., 2023). Additionally, auxin and BR share overlapping activities in the promotion of LR (Bao et al., 2004; Chaiwanon and Wang, 2015; Kim et al., 2007; Li et al., 2005; Mazzoni-putman et al., 2021) suggesting a possible positive role for BZR1 in upregulating targets important for callus formation. Given the vast difference in *BES1* and *BZR1* expression in CIM media and its correlation with the callus phenotypes of *bes1-D* and *bzr1-D*, we wanted to assess if a similar trend would be observed during the transition to SIM. We addressed this by transferring explants to SIM after a 21-day CIM treatment. In these conditions, and contrasting with what was observed for CIM, BES1-YFP signal increased dramatically within just 3 days of treatment in the newly formed SAM (Figure 3C). This is consistent with the phenotype displayed by *bes1-D* under SIM treatment, namely increased shoot formation (Figure 1). As for BZR1-YFP, the expression of this marker was reminiscent of that displaying CIM, *i.e.* high expression throughout the entire callus mass (Figure 3D). Contrary to our findings, it has been shown that BES1-GFP accumulates after callus induction, however those observations were based on callus induction from hypocotyl explants (Wu et al., 2022) and the discrepancies in the results could be a consequence of tissue specific roles of BES1 and possibly BZR1. Interestingly and consistent with our results, the authors also showed increased BES1-GFP signal after growth on SIM . Altogether, our data suggest that during CIM treatment, stabilization of BZR1 and consequent increased nuclear localization facilitates pluripotency acquisition and cellular proliferation, leading to increased callus production, while BES1 shows the opposite response. However, on SIM, both transcription factors appear to play important roles, as both are highly expressed upon subculturing of explants in this media.

**Figure 3.**
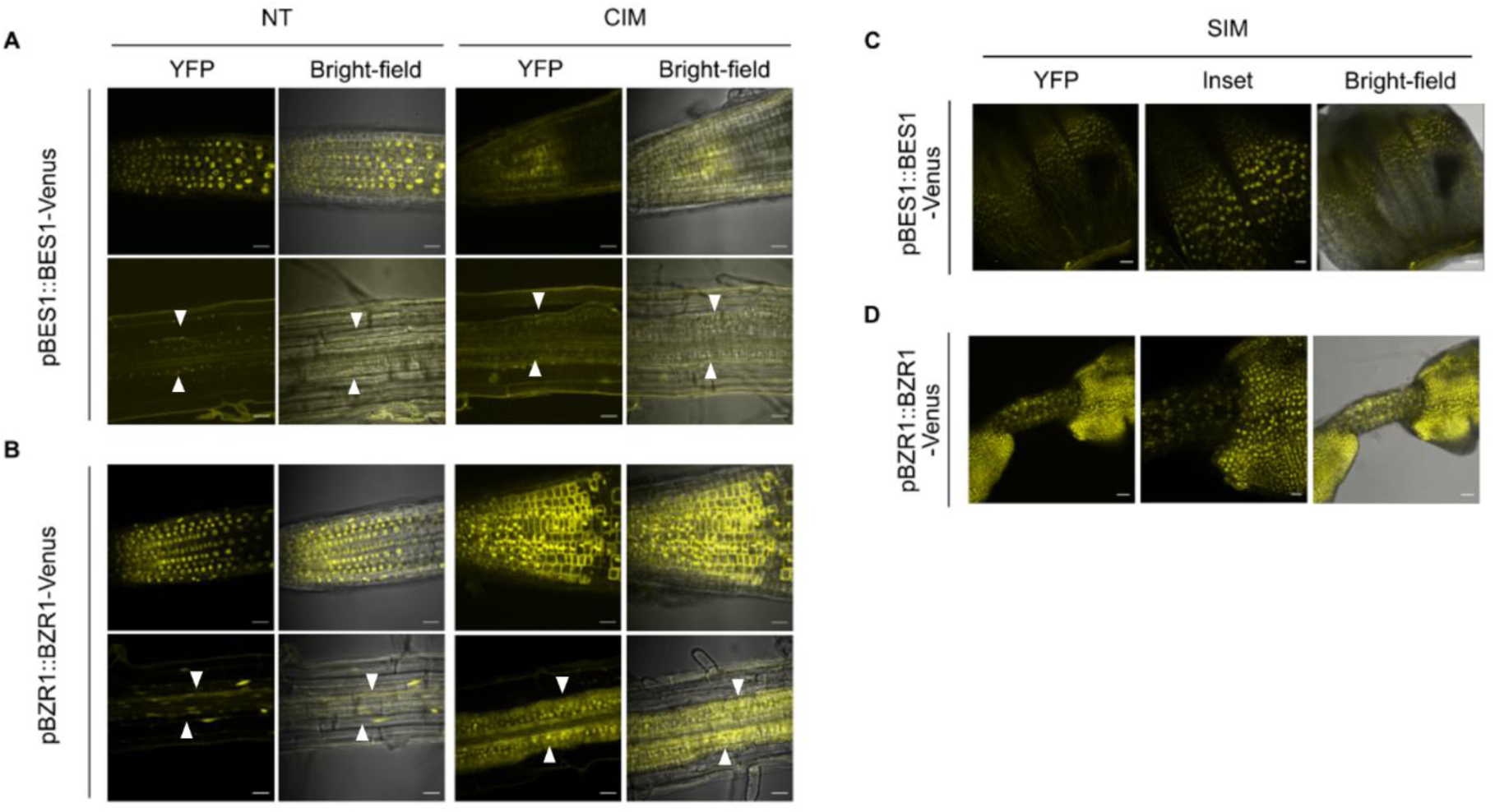
BES1 and BZR1 expression during callus formation and de novo shoot formation. A-B) Confocal images of root tips (top panel) and mature root tissue (bottom panel) in Col-0 background. a) pBES1::BES1-Venus and b) pBZR1::BZR1-Venus in 7-day-old root explants followed with 6-Day treatment of liquid ½ MS plus solvents or liquid CIM. n= 9 per condition. C-D) Confocal images of callus treated for 21-day CIM prior to 3 day SIM expressing c) pBES1::BES1-Venus and d) pBZR1::BZR1-Venus. n= 9 per condition. White arrows mark the XPP surrounding cells

### BZR1 binds and regulates ARF7 and ARF19 under callus formation

Our combined results showing enhanced *BZR1* expression on CIM and increased callus mass of *bzr1-D* suggest that BZR1 might have a role in mediating pluripotency acquisition. Given that BZR1 is known to bind and directly regulate the central pluripotency regulator *ARF7* (Zhou et al., 2013), we decided to examine the relative expression levels of *ARF7* and also *ARF19*, which often works redundantly to *ARF7*, during callus formation (Wilmoth et al., 2005) (Figure 4A-B). Consistent with our reasoning, expression of both *ARF*s was significantly elevated upon 21 days of CIM treatment in *bzr1-D* when compared to WT (Figure 4A-B). Encouraged by this result, we assessed whether BZR1 binds *ARF7* and *ARF19* promoter regions during callus induction through chromatin immunoprecipitation (ChIP). As indicated in Figure 4C, the promoter regions and initial gene segments of both *ARF7* and *ARF19* exhibit the presence of E-box and BRRE elements. Following the 21-day CIM treatment, we observed enriched binding of BZR1 to the promoter regions of *ARF7* and *ARF19* specifically (Figure 4D & figure S2). These results indicate that BZR1 not only regulates differential growth through *ARF7* in the hypocotyl (Zhou et al., 2013) but also during callus formation (Figure 4), further expanding the complex role of *BZR1* during developmental regulation (Chaiwanon and Wang, 2015; Gendron et al., 2012). Together these findings demonstrate that the binding of BZR1 leads to an increased expression of *ARF7* and *ARF19*, potentially facilitating callus formation.

**Figure 4.**
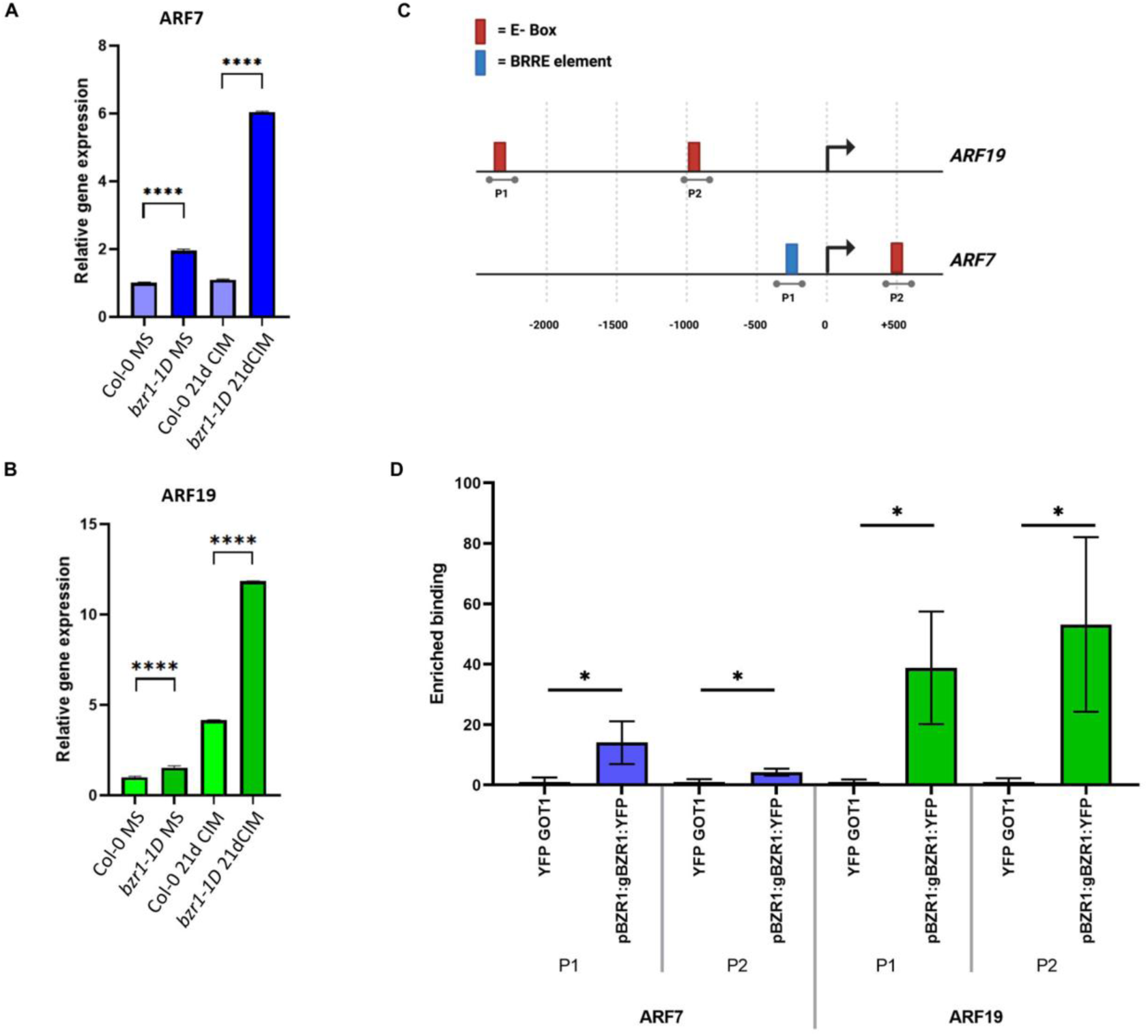
Callus induction enhances BZR1 binding to the promoters ARF7 and ARF19. A-B) Relative expression of A) ARF7 and B) ARF19 were analyzed in WT and bzr1-D. Expression was normalized to Actin 2 and is relative to Col-0 MS; i.e. root explants from 7-day old seedlings grown on MS media. Tissue was obtained from NT and 21-day CIM-treated root explants. Experiments were repeated 3 times and similar results were obtained. Bars indicate the SD of the mean. Significance was calculated via student t-test. Asterisks represent the significant difference from WT from each treatment, (***P<0.001). Representative figure is shown. Experiments were performed in biological triplicates with similar results C) Schematic overview over promoter regions for ARF7 and ARF19. E-box and BRRE-element are marked accordingly, and primer sets are boarding the sites. D) The DNA binding of pBZR1::BZR1-YFPto ARF7 and ARF19 promoter region was determined through quantitative RT-PCR. Fold change was calculated by normalizing to the YFP::GOT1 control for each sample . Error bars show SD from 3 biological replicates. Significance was calculated via student t-test (P<0,05). Asterisks represent the significant difference from YFP::GOT1 (control) (*P<0.1).

### bzr1-D enhanced callus formation is dependent on ARF7 and ARF19

In light of our results showing that BZR1 directly regulates *ARF7* and *ARF19* during CIM treatment, we wondered if the enhanced callus formation phenotype of *bzr1-D* was dependent on these two ARFs. To test this, we generated a triple mutant line, *bzr1-D arf7-1 arf19-2* (Figure 5), which showed that, consistent with our hypothesis, *bzr1-D* callus phenotype was abolished in the *arf7-1 arf19-2* background (Figure 5C & D). *bzr1-D/arf7-1/arf19-2* lines produced above-ground phenotype similar to *arf7-1/arf19-2 (Figure S3).* Previous studies have linked BIN2 to the regulation of LR organogenesis through direct phosphorylation of ARF7 and ARF19 (Cho et al., 2014) and surprisingly the BIN2 gain of function mutant, *bin2-1*, exhibited enhanced callus formation (Lee and Seo, 2017) similar to what we observed in *bzr1*-D (Figure 1). While it might seem counterintuitive that both *BIN2* and *BZR1* gain-of-function mutants could display similar phenotypes, one should note that *bin2-1* callus phenotype seems to be, at least in part, independent from brassinosteroid signalling, as supplementation with BL lead to further increase in callus formation of *bin2-1*. It is therefor possible that under CIM treatment BIN2 does not antagonize BZR1, and both end up promoting callus formation via *ARF7* and *ARF19*. Precisely, disruption of *ARF7* and *ARF19* in both *bzr1-D* (Figure 5) and *bin2-1* (Lee and Seo, 2017) backgrounds, led to the suppression of callus formation indicating that both BIN2 and BZR1 mediated callus formation is dependent on *ARF7* and *ARF19*. As LR initiation and callus formation share a developmental program, we wanted to evaluate LR phenotype of *bzr1-D* and its potential dependency on *ARF7* and *ARF19*. Interestingly and as seen in figure 5A & B, *bzr1-D* produced significantly more LRs than WT which was yet unreported for this mutant and in contrast to previous results showing no difference between Col-0 and *bzr1-D* (Cho et al., 2014). In agreement with our callus phenotype *bzr1-D arf7-1 arf19-2* was also unable to form LRs, a phenotype reminiscent of *arf7-1 arf19-2*, which altogether indicates that *bzr1-D* enhanced pluripotency arises from direct interaction and regulation of *ARF7* and *ARF19*.

**Figure 5.**
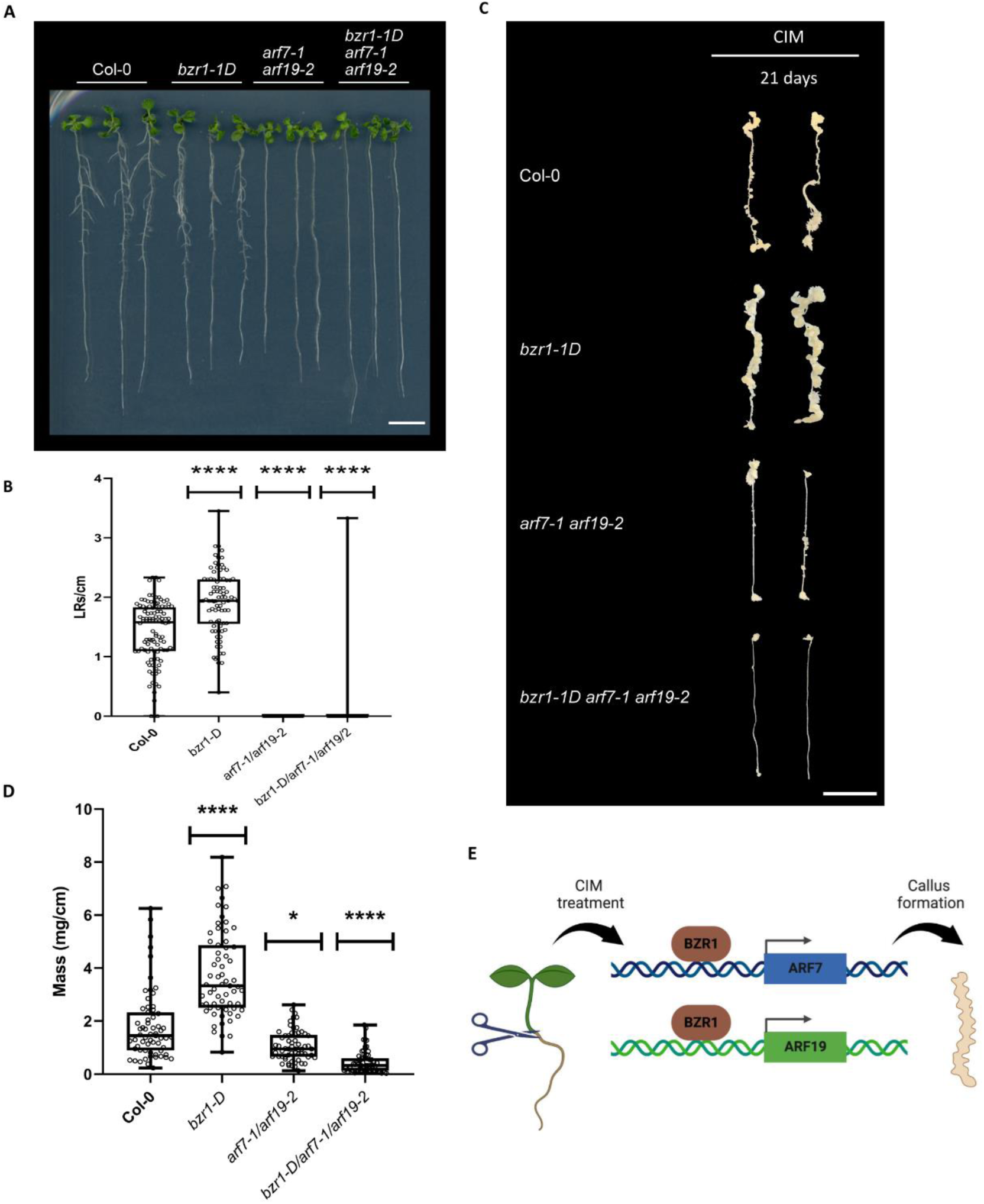
Enhanced callus and LR formation on bzr1-d are dependent on ARF7 and ARF19. A) Representative images of 10-day-old seedlings WT, bzr1-D, arf7-1 arf19-2 and bzr1-D arf7-1 arf19-2 on ½ MS media. Scale bar 1 cm B) Lateral root count of 10-day-old seedlings, WT, bzr1-1D. Significance was calculated via Kruskal Wallis non-parametric test. Asterisk, represents the significant difference from Col-0 (****P<0.0001). C) Representative images of callus forming root explants WT, bzr1-D, arf7-1 arf19-2 and bzr1-D arf7-1 arf19-2 from 21-day CIM, scalebar = 1 cm. D) Fresh weight of 21 -day root explants on CIM. Significance was calculated via Kruskal-Wallis non-parametric test (P<0,05). Asterisks represent the significant difference from WT Boxplots: centerlines show the median; the box limits the 25^th^ and 75^th^ percentiles; whiskers extend to the minimum and maximum. Experiments were made in three biological replicates with total n=36-106 (B), n=63 (D) . E) Graphical abstract; Upon CIM treatment BZR1 binds and directly regulate ARF7 and ARF19 to enhance callus formation. The figure was drawn using BioRender (https://www.biorender.com).

## Conclusion

In summary, our findings add additional knowledge to the comprehensive interplay between BR and auxin signaling, specifically BZR1, and its role during pluripotency acquisition. As others previously have shown, the two hormones promote and repress each other to accommodate the changes needed to obtain “normal” root growth (Ackerman-Lavert et al., 2021; Bao et al., 2004; Chaiwanon and Wang, 2015; Rovere et al., 2022). Here we put BZR1 as a direct contributor to LR formation and demonstrate an increased expression of *BZR1* in the dividing tissue during callus induction. Through direct regulation, BZR1 binds and activates the expression of key LR regulators *ARF7* and *ARF19*, essential for callus formation. Additionally, we propose a possible association with BZR1 and BES1’s effect in activating *de novo* organogenesis in root explants. In a recent study, BES1/BZR1 family of TF was shown to represses shoot formation under cell fate transition but this observation was based on studying hypocotyl explants (Wu et al., 2022). Similarly, BIN2 was shown to be a positive regulator of callus formation in studies based on observations of the root/shoot junction only (Lee and Seo, 2017). In both cases, no quantification of root explants was made. These apparent contradictions could be explained by the ability of BR, and consequently BZR1 and BES1, to both promote and repress different processes, exhibiting a dual regulatory function (Ackerman-Lavert et al., 2021). This could happen in a tissue-specific manner, which would point towards a model where BZR1, BES1 and possibly BIN2 can act as both repressors and promoters of callus formation and organogenesis dependent on the type of explant tissue. While our data provide evidence that excessive BZR1 promotes callus formation through regulation of *ARF7* and *ARF19*, more experiments are needed to fully comprehend the complicated process of explants forced into cell fate transition.

## Acknowledgement

We thanks Ana Caño-Delgado, Crag Genomica, for the *BES1-*D seeds, Wenqiang Tang, Hebei Normal University, for the *pBZR1::BZR1-YFP* seeds, and Jia Li, School of Life Science for the *pBES1::BES1-YFP* seeds. Confocal microscopy imaging was performed using equipment from the Center for Advanced Bioimaging (CAB) Denmark, University of Copenhagen. This work was funded by the Danish Research Agency grant to E.R. (DFF1-1032-00249B) and the Novo Nordisk Fonden (NNF190C0055222) to M.P..

## Methods and materials

### Plant growth and phenotypic characterization

For growth of Arabidopsis seedlings, ½ MS media (0.8% agar, 1% sucrose at 5.7 pH) was used, exposing the plants to a photoperiod of 16 hours (120 µE/m^2^/s), at 21 C°). Seed sterilization was accomplished with 1.3% bleach for 2 min, followed by 70% ethanol for 1 min, and washed three times with sterile water. We used following lines in this study: Col-0, *bzr1-D* (Wang et al., 2002), *bes1-D* (Ibañes et al., 2009), *arf7-1 arf19-2*(NASC), *bzr1-D arf7-1 arf19-2* (crossed), *pBZR1::BZR1-YFP* (W. Chen et al., 2019), *pBES1::BES1-YFP* (W. Chen et al., 2019), *YFP-GOT1* (YFP fusion to GOT1 (At3g03180), which is a protein involved in vesicle trafficking between Golgi and ER) (Geldner et al., 2009), *bzr1-c1* (constructed), *bzr1-c2*(constructed) and *bes1-ko* (salk_098634).

For experiments, 7-day-old seedlings were grown on solid half-strength MS before callus cut. For callus growth, explants were put on solid CIM media (3,1 g/L Gamborg’s B5 salts with vitamins, 20g/L Glucose, 0.5 g/L MES, 8g/L Agar, pH adjusted to 5.7 with 1 M KOH) containing 2,4-D (0.5mg/L) and kinetin (0.05mg/L). Incubation time was 21 days before moving them to solid SIM media (4.3 g/L Murashige and Skoog, 30 g/L sucrose, 8 g/L Agar, pH adjusted to 5,8 with 1M NaOH) containing BAP (1.0 mg/L) and IAA (0.1 mg/L). Images of callus were taken after 21 days on CIM and after 21 days on SIM.

The *bzr1-D/arf7-1/arf19-2* crosses were made and genotyped using Salk primer LbB1.3 and gene specific primers for *arf7-1* and *arf19-2*, and decapping primers for *bzr1-D*. After digesting with the NcoI (Thermo Scientific) restriction enzyme, a 25 bp band shift was observed in the *bzr1-D* mutant. Confocal Images of root explants were taken post 6-day treatment with either liquid half-strength MS or liquid CIM media. Confocal microscopy images were taken with LSM700 Zeiss confocal microscopy using 20x objective. Images were analyzed using ZEN2012 (Zeiss) software.

### RNA extraction and qPCR

Total RNA was extracted from samples as previously described in (Zuo et al., 2022) through TRIzol reagent (Thermo Scientific), treated with DNase (Thermo Scientific) and cDNA was reverse transcribed through Reverse Aid kit (Thermo Scientific) following the manufacturer descriptions. For qPCR Fast Sybr green mastermix (Thermo Fisher) was used and the reaction was performed on Quantstudio 1 and Quantstudio 5 (Applied Biosystems). Actin2 (AT3G18780) was used as an internal control. Primers used are in Supplementary table 1.

### Chromatin immunoprecipitation (ChIP)

The ChIP was carried out using the Chromotek Chromatin Immunoprecipitation protocol for *A. thaliana*(https://www.ptglab.com/products/pictures/pdf/ChIP_Apps_Note_GFP-Trap_for_A_thaliana.pdf). Approximately 2 g of fresh weight material per genotype was used, and samples were crosslinked in vivo for 15 min with formaldehyde (1%). After sonication (500-1000pb fragments), DNA concentration was measured and all the samples were diluted to the same DNA concentration. 20uL (ChromoTek) GFP beads were added and samples were incubated overnight at 4°C with gently agitation. De-crosslinked samples were then purified and used for real-time quantitative PCR. All ChIP-qPCR were repeated three independent experiments. Primers are listed in supplementary table 1. Enrichment was calculated relative to the input and expressed as fold change to the YFP GOT1 which was arbitrarily set to 1

### Crispr Cas9 construction

SgRNA was designed with the help of http://crispr.hzau.edu.cn/CRISPR2/ targeting the 5’ end of AT1G75080. Two primer sets were used. The primer pair was annealed followed by ligation using T4 ligase (Thermo Scientific) into pHEE401(pre-digested with bsa1) before transformation into *E. coli*. Sequencing was done before transformation sgRNA into GV3101. Construct was moved to Agrobacterium followed by transformation into plants by flower dipping method. Genotyping primers and sgRNA primers are listed in Supplemental table 1.

**Supplementary table 1.**
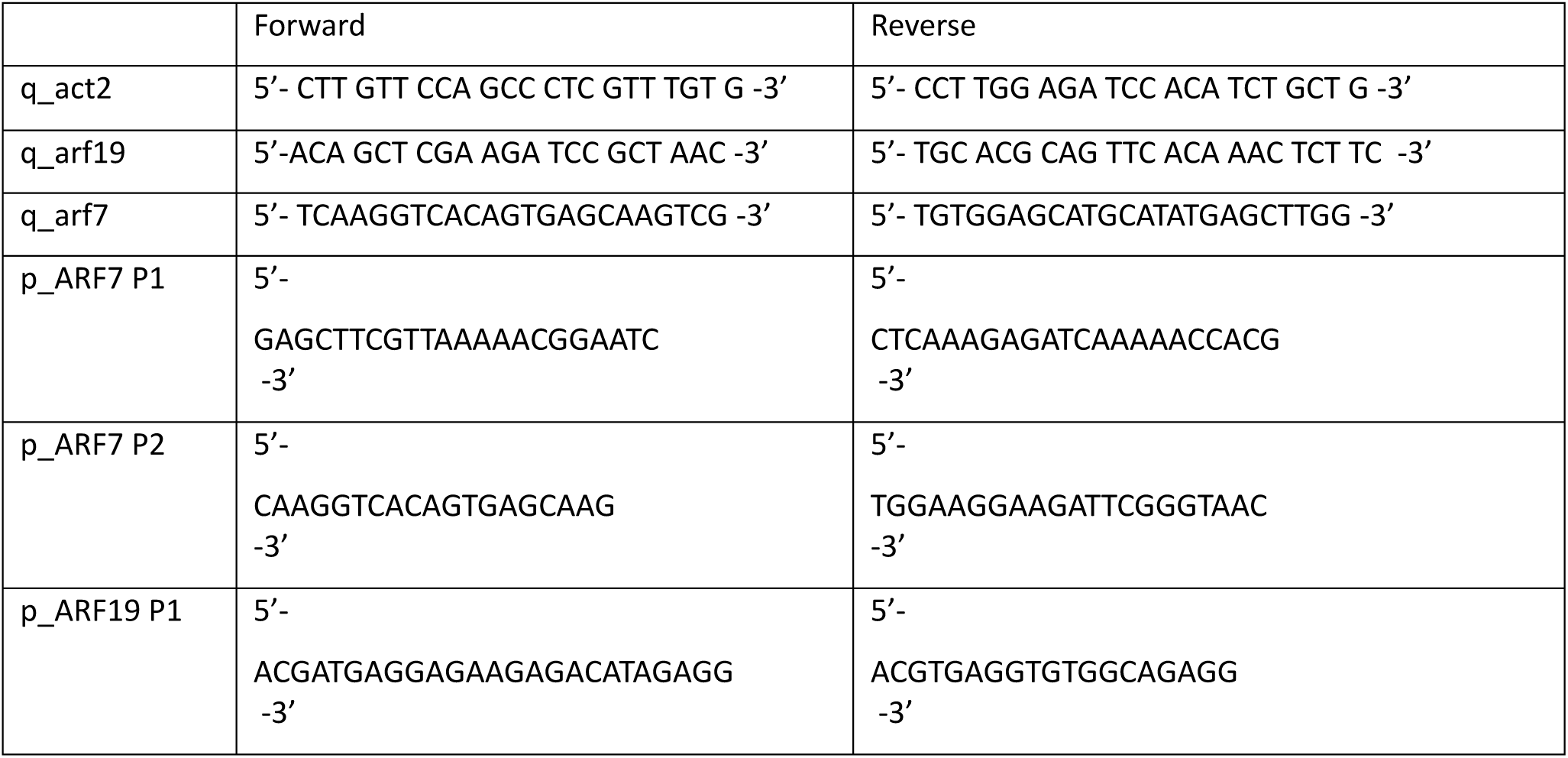
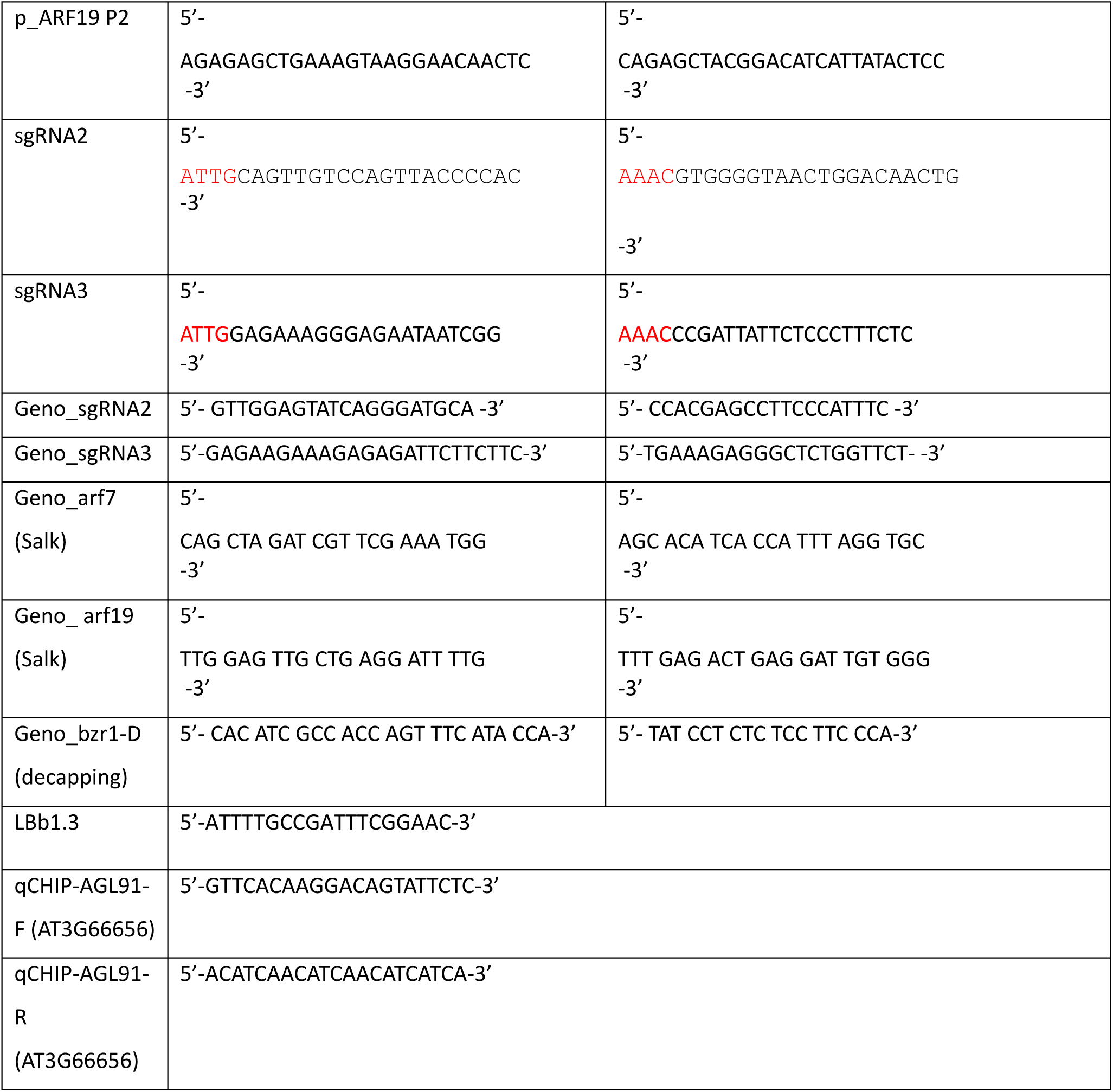
Primer list.

